# Single Cell Spatial Transcriptomic Profiling Identifies a LINE1 Associated Disarrayed Immune Microenvironment in Hepatocellular Carcinoma

**DOI:** 10.1101/2023.12.04.570014

**Authors:** Avril K. Coley, Cole Nawrocki, Bidish K. Patel, Amaya Pankaj, Zhoubo Guo, Matthew J. Emmett, Evan R. Lang, Yuhui Song, Katherine H. Xu, Cristina R. Ferrone, Vikram Deshpande, Linda T. Nieman, Martin J. Aryee, Joseph W. Franses, David T. Ting

**Affiliations:** Department of Surgery, Massachusetts General Hospital Harvard Medical School; Boston, MA, USA; Mass General Cancer Center, Harvard Medical School; Charlestown, MA, USA; Department of Medicine, Massachusetts General Hospital, Harvard Medical School; Boston, MA, USA; Section of Hematology-Oncology, Department of Medicine, University of Chicago; Chicago, IL, USA; Department of Surgery, Cedars-Sinai Hospital; Los Angeles, CA, USA; Department of Pathology, Beth Israel Deaconess Medical Center, Harvard Medical School; Boston, MA, USA; Department of Biostatistics, Harvard T.H. Chan School of Public Health; Boston, MA, USA; Department of Data Sciences, Dana-Farber Cancer Institute; Boston, MA, USA; Broad Institute of Harvard and MIT; Cambridge, MA, USA

**Keywords:** Spatial transcriptomics, hepatocellular carcinoma, tumor microenvironment, immune microenvironment, LINE1, repeat RNAs

## Abstract

**Purpose:** Hepatocellular carcinoma (HCC) is a lethal malignancy driven by complex interactions between cancer cells, immune cells, and additional stromal cells in the tumor microenvironment (TME). The LINE1 retrotransposon is a ubiquitous repeat RNA whose de-repression leads to significant cancer cell-intrinsic and TME changes that promote aggressive tumor characteristics. We leveraged single cell spatial transcriptomic profiling to characterize the relationship between LINE1 and differences in the heterogeneous HCC TME.

**Experimental Design:** We applied our profiling methodology to a cohort of 23 tissue specimens collected from patients who had undergone liver resection or transplantation and validated it in a partially-overlapping similar cohort of 39 specimens using RNA in-situ hybridization (RNA-ISH).

**Results:** We found that LINE1-high tumors and LINE1-high single HCC cells exhibited a de-differentiated, stem-like, and inflammatory phenotype. Furthermore, within individual tumors, LINE1 high cancer cells associated spatially with one another and excluded the larger, organized immune cell conglomerates seen in LINE1 low tumors. Finally, we found that LINE1 RNA expression correlated with worse overall survival in the larger expanded retrospective cohort.

**Conclusions:** Our study is the first to show a clearly disorganized immune TME in HCC driven by LINE1 expression, and this observation correlated with poor survival for patients whose tumors expressed large amounts of the LINE1 repeat RNA. These results provide further evidence of how effective anti-tumor immune responses contribute to cures after definitive surgery and may lead to novel biomarkers or drug targets in HCC.

**TRANSLATIONAL RELEVANCE:** The viral-like LINE1 retrotransposon is known to influence tumor cell state and the immune response in a variety of cancer. Here, we have used single cell spatial transcriptomic profiling to resolve repeat and coding gene RNA expression in a cohort of hepatocellular carcinoma (HCC) patients. LINE1 RNA expression in HCC tumor cells was correlated with an undifferentiated stem-like cancer state and a disorganized, sparse immune infiltrate. Using in situ hybridization on an expanded validation cohort, we noted significantly worsened survival in the LINE1 high group. Altogether, LINE1 repeat RNA is a tumor intrinsic biomarker of more aggressive features that can be used for risk stratification and a potential biomarker for response to immunotherapies that merits further investigation.

## INTRODUCTION

Hepatocellular carcinoma (HCC) is a leading cause of global cancer morbidity and mortality (1). Modern medical treatment regimens for advanced stage HCC have improved significantly over the past several years, with the emergence of multiple therapies inhibiting either pathologically activated vasculature, dysregulated immune checkpoints, or combinations thereof (2). These features have highlighted the importance of therapeutically targeting the HCC tumor microenvironment (TME). However, only a minority of patients derive significant clinical benefit from these regimens (3–5), and most patients eventually succumb to their disease (6,7). These factors highlight the importance of an improved understanding of the dynamic cellular interactions in the TME of HCC tumors, which has the potential to broaden our understanding of existing drug mechanisms and motivate the subsequent development of novel drug targets and biomarkers.

Our group and others have characterized the relationship of repeat RNA expression with the TME across cancers (8–15). These relationships appear to be linked to differences in repeat RNA sequence (16) and secondary structure (17), which both contribute to the activation of innate immune responses (17–26). Prior work has demonstrated the worsened prognosis of HCC patients with hypomethylation of the LINE1 repeat RNA (27–30) and evidence of retrotransposition events in viral and non-viral HCC (31). Further, there is mounting evidence of LINE1 retrotransposition events in viral and non-viral interplay between HBV and HCV driven HCC oncogenesis with LINE1 (32,33). More generally, expression of specific repeat RNA species tends to correlate with distinct patterns of tumor immune infiltrates (8,9,15).

Given the known viral drivers of cirrhosis and HCC (34), we hypothesized that endogenous repeat viral-like elements are important in HCC carcinogenesis and immunological response. To fully characterize the relationship of repeat elements with tumor cells and the surrounding microenvironment, we utilized high-plex spatial tissue profiling technologies. These technologies have many of the benefits of sequencing-based and proteomic cell profiling methodologies while retaining the spatial information inherent to conventional tissue profiling technologies such as in situ hybridization or immunohistochemistry (35). Our previous work in HCC utilized a regional whole transcriptome analysis of tumor cells, endothelial cells, and immune cells revealing improved resolution of tumor cell subtyping and initial ligand-receptor signaling between tumor cells and the vasculature (36).

In this study, we have expanded on these previous studies by utilizing spatial molecular imaging of 957 genes (including several repeat RNA species) in HCC tumor resection samples to understand the relationship of repeat RNAs to tumor cell states and the microenvironment response. This is the first integrated dataset characterizing repeat RNA distributions and single cell spatial architecture in a cohort of samples derived from human HCC resection or transplantation specimens. We found that the HCC LINE1-high tumor cells are enriched for a stem cell phenotype with loss of hepatocyte differentiation markers. LINE1-high HCC tumors had a more disorganized immune response compared to LINE1-low HCC tumors that had regional hubs of diverse immune cell infiltrates similar to tertiary lymphoid structures. Using RNA in situ hybridization (RNA-ISH) for LINE1, we found that LINE1-high compared to LINE1-low HCC tumors had significantly worsened survival after surgical resection or transplantation. Taken together, these findings support the importance of LINE1 and other repeat RNAs in HCC development and as prognostic markers of aggressive behavior.

## MATERIALS AND METHODS

### Ethics statement, tissue procurement and annotation

All patient-oriented research was conducted in accordance with both the Declarations of Helsinki and Istanbul. Patient tumor materials were obtained under Massachusetts General Hospital IRB protocol 2011P001236 and Dana-Farber Harvard Cancer Center IRB protocol 02-240. Archived FFPE samples in tissue microarray (TMA) format from who underwent surgical resection or liver transplantation for the treatment of hepatocellular carcinoma between March 2004 and December 2015 were obtained. Clinical and pathologic data was obtained through review of the Massachusetts General Hospital electronic medical record. There is no blinding, randomization, or power analysis relevant for this study.

### Spatial transcriptomics data acquisition

A total of 23 patient FFPE HCC tumors on TMAs were evaluated with spatial molecular imaging (SMI) following published standard protocol (37). In brief, 5-µm FFPE sections were baked overnight at 60°C to ensure section adherence to the glass slides. Then the baked samples went through deparaffinization, proteinase K digestion, and heat-induced epitope retrieval (HIER) procedures to expose target RNAs and epitopes using Leica Bond Rx system. After rinsing samples with DEPC H_2_O twice, samples were incubated in 1:1000 diluted fiducials (Bangs Laboratory) in 2X SSCT (2X SSC, 0.1% Tween 20) solution for 5 min at room temperature. Excessive fiducials were removed by rinsing the samples with 1X PBS, followed by fixation with 10% neutral buffered formalin (NBF) for 5 min at room temperature. Fixed samples were rinsed with Tris-glycine buffer (0.1 M glycine, 0.1M Tris-base in DEPC H_2_O) and 1X PBS for 5 min each before blocked using 100 mM N-succinimidyl acetate (NHS-acetate, ThermoFisher) in NHS-acetate buffer (0.1 M NaP, 0.1% Tween 20, pH 8 in DEPC H_2_O) for 15 min at room temperature. Prepared samples were rinsed with 2X saline sodium citrate (SSC) for 5 min and then Adhesive SecureSeal^TM^ Hybridization Chamber (Grace Bio-Labs) was placed to cover the sample. ISH probe mix (1 nM ISH probes, 1X Buffer R, 0.1 U/µL SUPERase●In™ in DEPC H_2_O) was prepared by denaturing 980-plex RNA ISH probes at 95°C for 2 min and then placed on ice before making ISH probe mix. Hybridization occurred at 37°C overnight after sealing the chamber to prevent evaporation. After the overnight hybridization, samples were washed with 50% formamide (VWR) in 2X SSC at 37°C for 25 min for 2 times, rinsed with 2XSSC for 2 min for 2 times at room temperature, and then blocked with 100 mM NHS-acetate for 15 min. After blocking, the samples were washed twice using 2X SSC for 2 min at room temperature. A custom-made slide cover was attached to the sample slide to form a flow cell. Prepared samples were loaded to the CosMx SMI instrument and went through data collection, image processing, feature extraction, and cell segmentation procedures following published protocols (37). Transcript profiles of individual cell were generated by combining target transcript location and cell segmentation information and then fed into downstream data analysis (see Supplement for more details).

### RNA *in situ* hybridization (RNA-ISH)

A total of 39 patient FFPE HCC tumor sections on TMAs were evaluated with RNA-ISH Automated chromogenic RNA-ISH assay was performed using the Advanced Cell Diagnostics (ACD) RNAscope 2.5 LS Duplex Reagent Kit - RED (Catalogue No. 322150) on the BondRx 6.0 platform (Leica Biosystems Inc., Buffalo Grove, IL). Assay was performed using custom probes from ACD in Channel 1 against HERVK (Catalogue No. 469838) at original concentration, HERVH (Catalogue No. 433558) at a 1:5 dilution, HSATII (Catalogue No. 504078) at a 1:7500 dilution, and LINE1 (Catalogue No. 565098) at a 1:50 dilution. 5 μm sections of FFPE embedded tumor microarrays were mounted on Fisherbrand Superfrost Plus glass slides, baked for 55 minutes at 60°C, and placed on the BOND RX for processing. The RNA unmasking conditions for the tissue consisted of a 15-minute incubation at 95°C in Bond Epitope Retrieval Solution 2 (Leica Biosystems) followed by 15-minute incubation with Proteinase K which was provided in the kit. Probe hybridization was done for 2 hours at 42C with RNAscope probes which were provided by ACD.The RNA-ISH slides were imaged with the Motic EasyScan Infinity Digital Pathology Scanner at 40x magnification. RNA quantification was performed with the Halo Image Analysis Platform by Indica Labs. Individual tissue areas on the TMA corresponding to each patient were annotated. In each tissue area, cellular segmentation was performed by detection of hematoxylin-stained nuclei and the RNA-ISH probe was detected by red chromogen. For HERVK, HERVH, and HSATII, the average number of probe copies per cell in the tissue area was quantified. For LINE1, due to the density of stain in the samples with the highest expression, accurate detection of individual red copies was challenging. LINE1 expression was therefore quantified as the probe detected per µm^2^ tissue area.

## RESULTS

### Single cell spatial transcriptomic profiling of HCC FFPE tissue

We analyzed FFPE TMA sections derived from 23 patient resection specimens using spatial molecular imaging (**Fig. 1A**). The probe set utilized for this experiment leveraged a 950 gene panel with additional custom “spiked-in” probes targeting repeat RNAs (HERVK, HSATII, LINE1-ORF1, LINE1-ORF2) as well as known HCC markers (*ASGR1*, *GPC3*, *LIN28B*) (**Table S1**). We utilized consecutive hematoxylin and eosin stained FFPE sections to guide the selection by an anatomic pathologist (BKP) of fields of view (FOVs) that contained representative areas of different TMA cores (**Fig. 1B, Fig. S1-2**). We then utilized immunofluorescent co-stains including a nuclear DAPI stain combined with fluorescent-conjugated primary antibodies targeting CD45, keratin 8/18 (CK8/18), pan-cytokeratin (panCK), and beta-2-microglobulin (B2M) to perform cell boundary segmentation (38) that is needed to identify and map each transcript to a specific cell (**Fig. 1C**). With this approach, after applying quality control filters to remove cells with adequate transcript density, we identified 158,971 cells with a per FOV median of 1081 cells (range 229-2812 cells) with an average of 244 transcripts/cell and 81 unique genes/cell (**Fig. 1D**).

**Figure 1.**
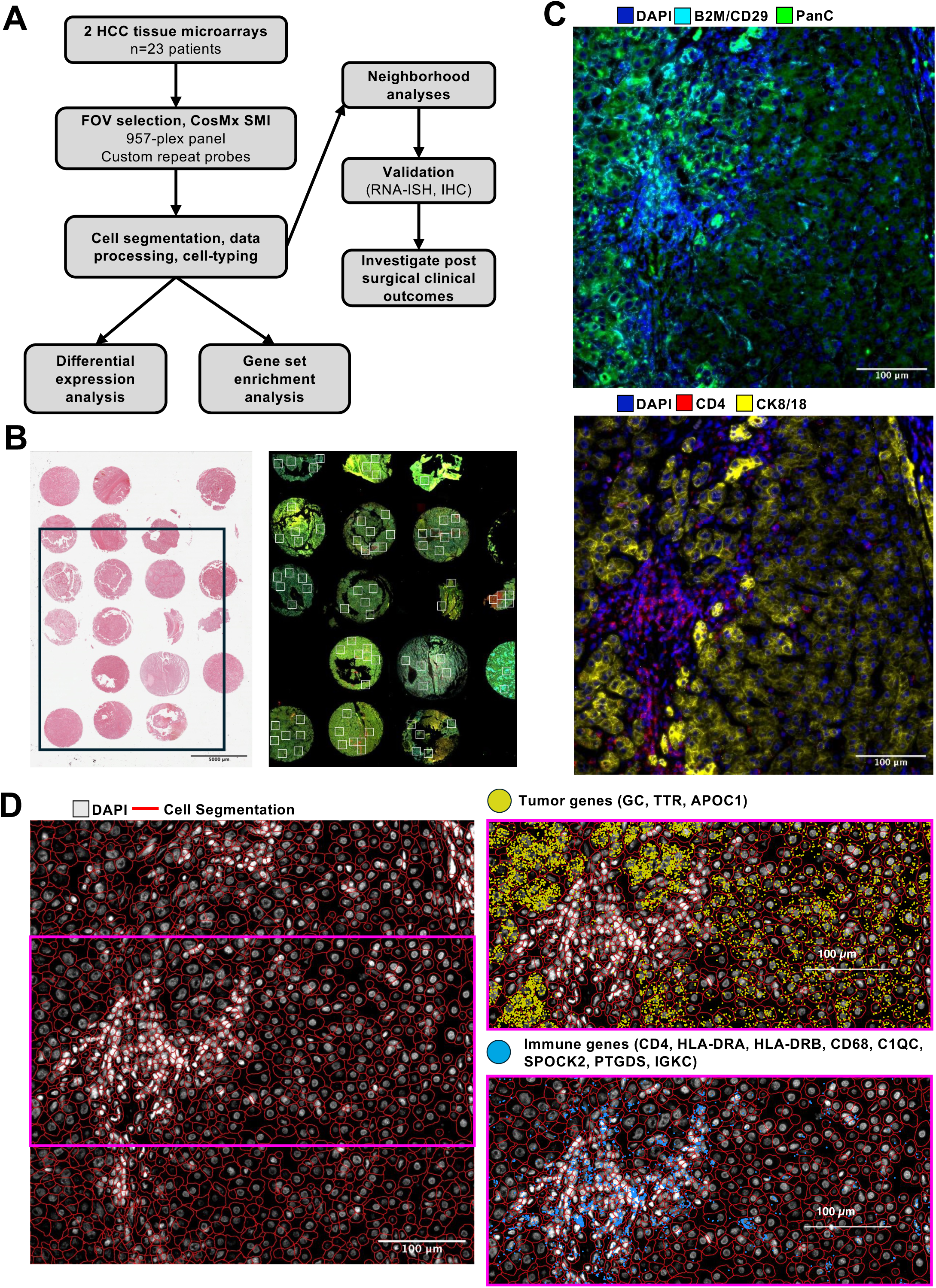
Overview of spatial molecular imaging (SMI) experiment utilizing HCC FFPE sections. (**A**) Experimental workflow of spatial transcriptomic profiling experiment. (**B**) Serial section H&E staining (left) and corresponding original immunofluorescence staining (right) for core tissue samples used in the SMI experiment from TMA2. For TMA1, see Figure S3. White rectangles depict fields of view (FOVs), within which SMI was conducted. Green: pan-cytokeratin, red: CD45, yellow: CK8/18, cyan: CD298/B2M. (**C**) Immunofluorescent staining used for cell segmentation in a representative FOV: TMA2, FOV60. (**D**) Cellpose cell segmentation results (left) and representation of SMI transcript detection (right) used to identify tumor and immune cells in the same FOV as in 1C.

Next, in combination with manual labeling, we utilized a combination of unsupervised and supervised techniques (39) to annotate cell identity (**Fig. 2A**). This resulted in classification of 96,677 tumor cells and 62,294 non-tumor cells (**Fig. 2B, Fig. S3**). For each cell type cluster, we found the top 1-5 genes for each cluster for which average expression was uniquely greater than in the other clusters to confirm cell identity (**Fig. 2C**) (40), and we further validated with differential expression analyses as performed in previous studies (**Fig. S4**) (41). We found that the HCC tissue cellular composition was consistent with those from prior dissociated HCC single cell datasets (42). We noted high variability in the composition of individual FOVs, with 25-90% of the cells in the HCC tumor FOVs had a range of 25-90% cancer cells, 10-30% stromal cells (fibroblasts, endothelial cells), and 10-50% immune cells (lymphoid cells, myeloid cells) (**Fig. 2D**). In summary, we utilized a subcellular resolution spatial transcriptomic technology to annotate and profile the in situ, spatially defined gene expression profiles of cells comprising systemic therapy-naïve HCC tumor microenvironment.

**Figure 2.**
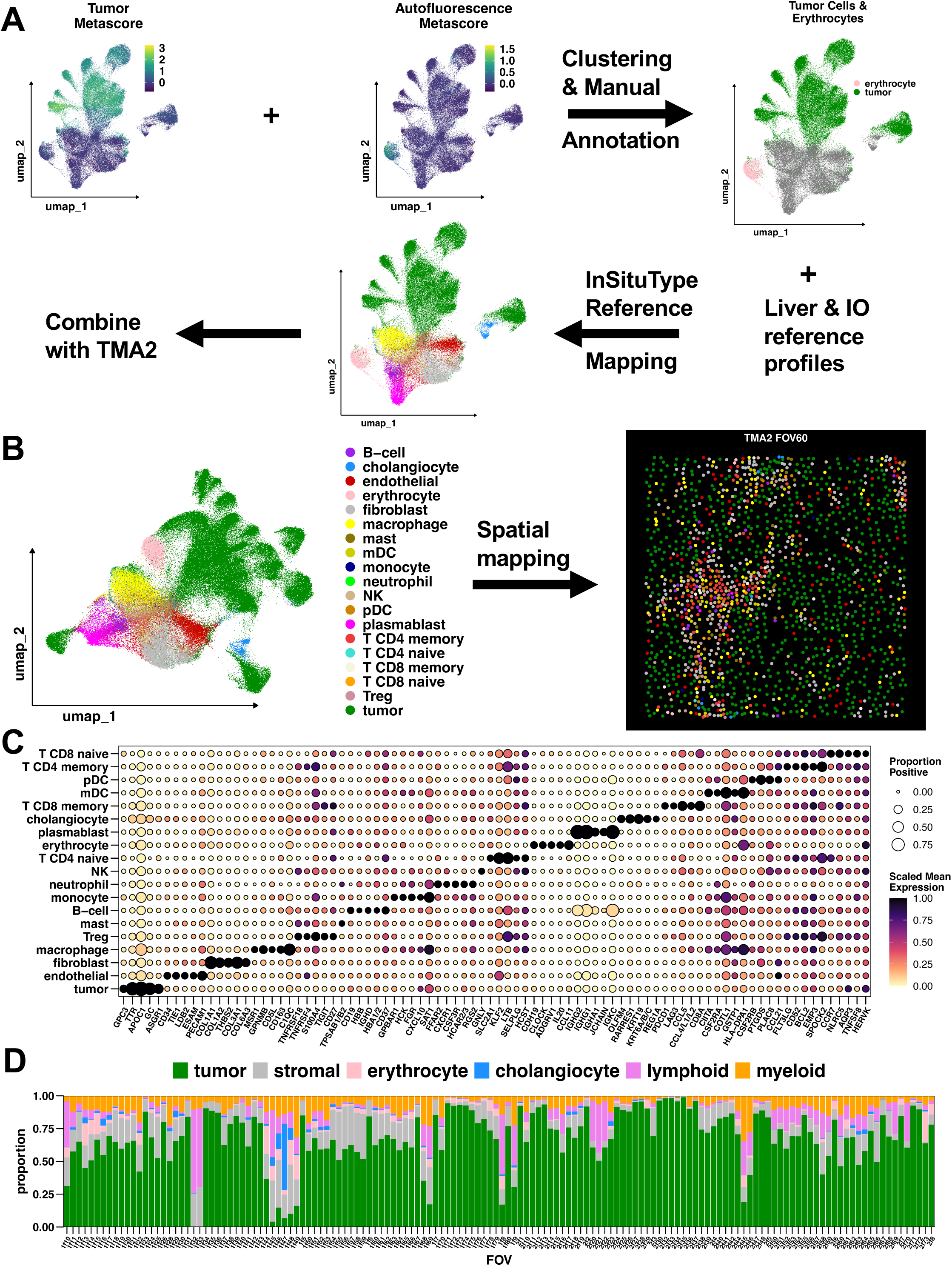
Subcellular spatial molecular imaging (SMI) accurately identifies cell types in HCC tumor microenvironment. (**A**) Schematic of cell-typing workflow, which included both manual annotation and supervised clustering, applied to TMA1. The same workflow was applied to TMA2 (see methods). (**B**) Final cell type annotations visualized on the UMAP embedding for the complete SMI dataset (left). Cells mapped back onto their physical locations in space, pseudo-colored by final cell type annotation for the same FOV in 1C (right). (**C**) Dot plot depicting expression of marker genes found algorithmically (see methods) for each of the identified cell type clusters. (**D**) Bar plot depicting the composition of each FOV in the complete SMI dataset in terms of major cell type categories.

### LINE1 expression correlates with aggressive cancer phenotypes and disorganized immune niches

Given prior work in other tumor types highlighting distinct immune niches associated with expression of various repeat RNAs, we then examined the spatial expression profiles of repeat RNA species using custom “spike in” oligonucleotide probes. Given prior literature highlighting the role of LINE1 methylation and retrotransposition in HCC (27–31,33), we focused our analysis on LINE1 ORF1 RNA (LINE1-ORF1) across the samples (**Fig. 3A**). Separating samples into the top tercile of LINE1-ORF1 (LINE1-high) and bottom two terciles (LINE1-low), we identified 579 differentially expressed genes, with 511 enriched in LINE1-high tumor samples and 68 enriched in LINE1-low tumor samples (**Fig. 3B**, **Table S2**). The gene set enriched in the LINE1-high subset included multiple additional repeat RNAs (LINE1-ORF2, HERVK, HSATII) and many additional genes typically associated with aggressive HCC behavior: WNT pathway ligands (*WNT3*, *WNT7A*, *WNT7B*, *WNT9A*) and receptors (*FZD1*, *FZD5*, *FZD7*, *FZD8*), stemness associated genes (LEFTY1, NDRG1, POU5F1), and multiple interferon pathway genes (*IFNL2/3*, *IFNGR2*) (**Fig. 3C**). The gene set enriched in the LINE1-low subset included multiple genes associated with normal hepatocyte function (*APOA1*, *APOC1*, *FGG*, *GLUL*).

**Figure 3.**
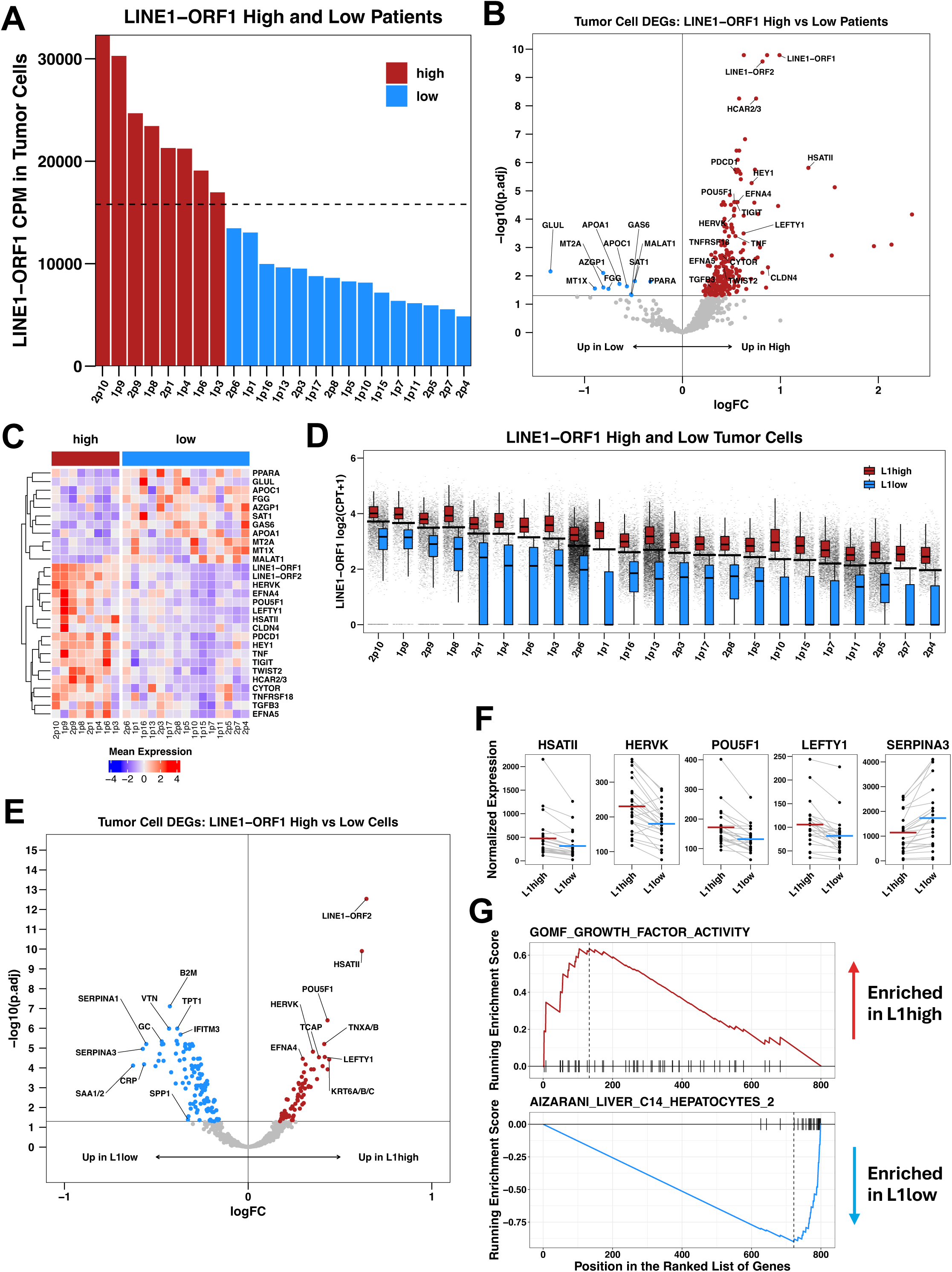
HCC expression of LINE1 is associated with a de-differentiated, stem-like state. (**A**) Bar plot depicting the definition of LINE1-ORF1 high and low patient groupings in the SMI dataset. The dashed line represents the upper tercile for net tumor cell LINE1-ORF1 counts per million (CPM) across all patients. (**B**) Volcano plot summarizing differential expression analysis comparing tumor cells in LINE1-ORF1 high patients and LINE1-ORF1 low patients, as defined in 3A. P-values are FDR-adjusted. Significance threshold = 0.05. (**C**) Heatmap depicting average normalized expression of selected differentially expressed genes identified in 3B among tumor cells in each patient. Values have been row scaled for visualization. (**D**) Box plots depicting the definition of LINE1-ORF1 high and low tumor cell groupings within each patient in the SMI dataset. The black crossbars represent the upper tercile for normalized LINE1-ORF1 expression within each patient. (**E**) Volcano plot summarizing differential expression analysis comparing LINE1-ORF1 high tumor cells and LINE1-ORF1 low tumor cells in a patient-paired manner. P-values are FDR-adjusted. Significance threshold = 0.05. LINE1-ORF1 not shown. (**F**) Patient-paired differences in total normalized expression of selected differentially expressed genes identified in 3E between LINE1-ORF1 high and low tumor cells. The crossbars represent means. (**G**) Gene set enrichment analysis plots generated from the results depicted in 3E.

Next, we examined the individual cell variability in LINE1 expression within each FOV (**Fig. 3D**). Similarly to the patient-level data, differential expression analysis (**Fig. 3E-F, Table S3**) showed enrichment of co-regulated repeat RNAs (HSATII, HERVK) and stemness related genes (*LEFTY1*, *POU5F1*) in the LINE1-high single cells. The LINE1-low cancer cells were enriched in markers of hepatocyte differentiation (*APOA1*, *FGG*, *SAA1/2, SERPINA3*). Gene set enrichment analysis of the LINE1-high versus LINE1-low cells confirmed a diminution in well-differentiated gene signatures (**Table S4**) and an enrichment of growth factor signaling (**Table S5**) changes for LINE1-high cells (**Fig. 3G**).

We then sought to understand the spatial structure of the TME niches associated with LINE1 expression, and we utilized spatial colocalization analysis to compare LINE1-high versus LINE1-low tumors. First, we noted, within each patient’s TME, that LINE1-high cells tended to cluster with one another, and LINE1-low cells also clustered with one another (**Fig. 4A-B**). Next, we noted several significant differences in the immune niches of patients with LINE1-high tumors compared to those of LINE1-low tumors. Cellular neighborhoods in the patients with low LINE1 tended to display large groups of co-localized immune cells, whereas neighborhoods in patienst with LINE1-low tumors had more dispersed immune cells (**Fig. 4C-D**). Relative to LINE1-low niches, LINE1-high niches featured fewer correlated CD4+ T cell – NK cell pairs and myeloid dendritic cell – endothelial and – monocyte pairs. Overall, the structure of the immune niche within LINE1-high cancer micro-regions showed overall statistically significantly lower immune co-localization (**Fig. 4C-E**).

**Figure 4.**
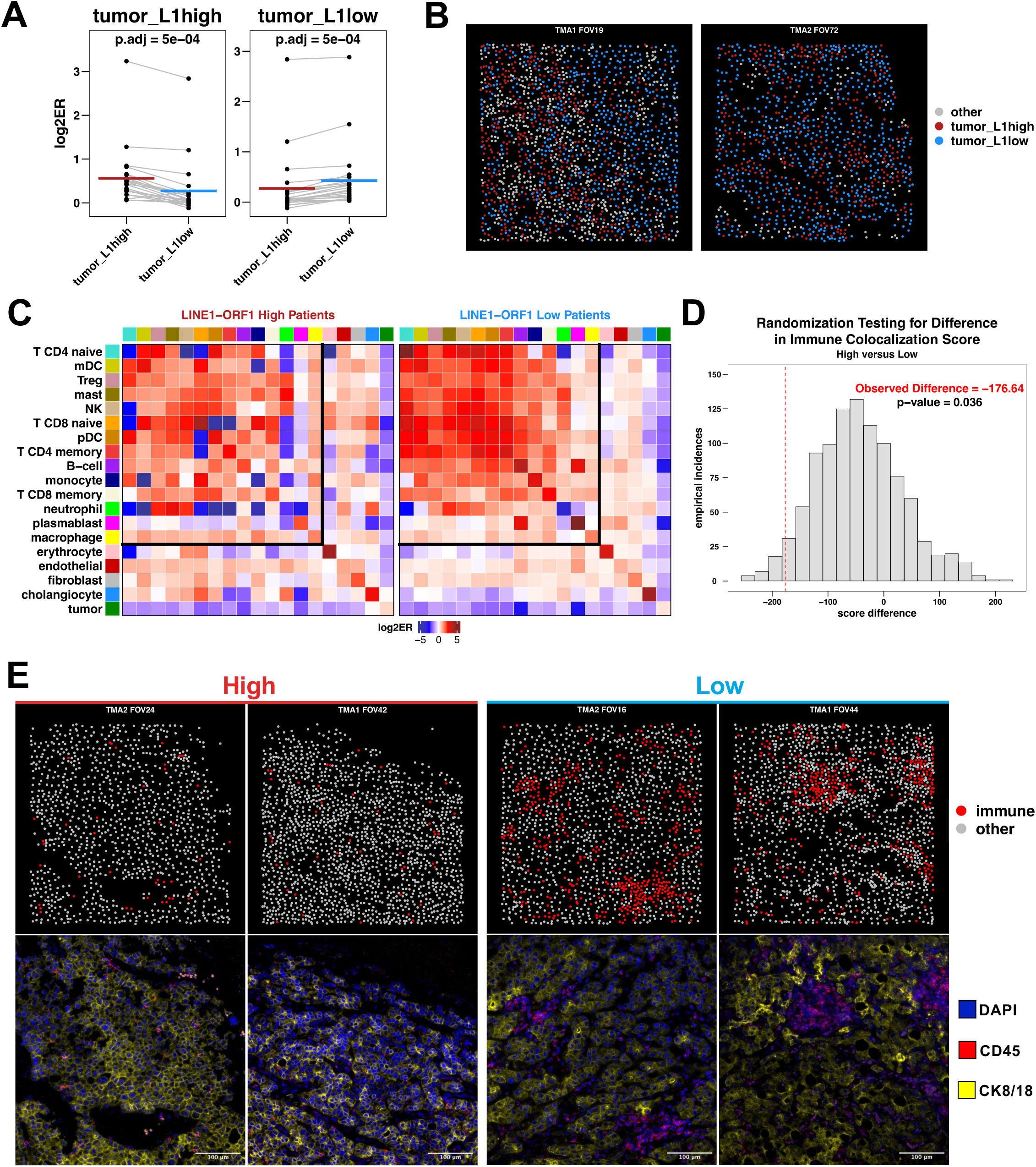
High LINE1 expression within HCC tissue leads to a disorganized, sparse immune microenvironment. (**A**) Patient-paired differences in LINE1-ORF1 high and low tumor cells’ colocalization patterns with themselves and one another. ER = enrichment ratio (see methods). P-values computed by paired Wilcoxon Rank Sum Test and FDR-adjusted. For other cell types, see Figure S6a. (**B**) Representative spatial maps for two FOVs depicting tumor niches comprised mainly of cells with the same LINE1-ORF1 group identity, as predicted by 4A. (**C**) Neighborhood enrichment analysis depicting pairwise colocalization for all cell type pairs in LINE1-ORF1 high patients (left) and LINE1-ORF1 low patients (right). (**D**) Randomization testing for difference in immune organization score between LINE1-ORF1 high and low patients. The red dashed line represents the observed difference which was compared to the shown empirical null distribution (see methods) to compute the reported p-value. (**E**) Representative immunofluorescent staining images and corresponding spatial maps for four FOVs illustrating differences in immune cell organization in LINE1-ORF1 high patients (left) and LINE1-ORF1 low patients (right).

Taken together, our results show that LINE1 expression within hepatocellular carcinoma cells correlates with expression of genes associated with aggressive cancer features and with a significantly disarrayed immune microenvironment. LINE1 expression levels may also be implicated in explaining how tumor cells from the same neighborhood arrange themselves. We next sought to validate and extend our spatial transcriptomic data using orthogonal techniques.

### Validation and clinical significance of LINE1 expression in HCC tissue

We validated repeat RNA expression using RNA in situ hybridization (RNA-ISH) across a cohort of 39 HCC tumor samples. Characteristics of the patient population from the expanded RNA-ISH cohort are summarized in **Table 1**. The median age of patients at the time of operation was 60 years old. Most patients had underlying cirrhosis (29/39; 74%) with the most common etiology being chronic HCV infection (15/39; 38%) followed by chronic HBV infection (6/39; 15%). A total of 30 patients underwent surgical resection (77%) and 9 patients had liver transplantation (23%). The five year overall survival proportion of this cohort was 51%. The demographic data in our cohorts are generally representative of published epidemiologic and outcome data (43).

**Table 1.** Patient characteristics.

|  |  | **Spatial** | **RNA-ISH** |
| --- | --- | --- | --- |
| **Sex (n, %)** | *male* | 15 (65%) | 28 (72%) |
|  | *female* | 8 (35%) | 11 (28%) |
| **Age (years)** | *median* | 63 | 60 |
|  | *range* | 45-82 | 39-85 |
| **Cirrhosis (n, %)** | *absent* | 15 (65%) | 10 (26%) |
|  | *present* | 8 (35%) | 29 (74%) |
| **Cirrhosis etiology (n, %)** | *alcohol* | 3 (13%) | 3 (8%) |
|  | *HBV* | 2 (9%) | 6 (15%) |
|  | *HCV* | 8 (35%) | 15 (36%) |
|  | *metabolic* | 2 (9%) | 3 (8%) |
|  | *other, NOS* | 8 (35%) | 12 (31%) |
| **Operation (n, %)** | *resection* | 20 (87%) | 30 (77%) |
|  | *transplant* | 3 (13%) | 9 (23%) |
| **Tumor size (cm)** | *median* | 4 | 4.3 |
|  | *range* | 1.5 - 13 | 1.2 - 15 |
| **Differentiation (n, %)** | *well* | 4 (17%) | 7 (18%) |
|  | *well-moderate* | 0 (0%) | 1 (3%) |
|  | *moderate* | 13 (57%) | 25 (64%) |
|  | *moderate-* | 3 (13%) |  |
|  | *poor* |  | 3 (8%) |
|  | *poor* | 3 (13%) | 3 (8%) |
| **Lymphovascular inv (n, %)** | *yes* | 11 (48%) | 16 (41%) |
|  | *no, NOS* | 12 (52%) | 23 (59%) |
| **Serum alpha-fetoprotein (ng/mL)** | *median* | 7.8 | 31 |
|  | *range* | 0 - 37510 | 0 - 37510 |

Digital imaging of RNA-ISH-stained TMA sections, followed by cellular segmentation and RNA-ISH signal quantification using the HALO AI platform (**Fig. S5B**), was used to quantify expression of HERVK, HERVH, HSATII, and LINE1 in the HCC tumor tissue (**Fig. 5A, Fig. S5C**). As for the spatial transcriptomic data, patient samples were divided into terciles based on the expression of each repeat RNA element. Patients with LINE1 expression in the upper tercile were designated as “high” and samples in the middle and lower terciles were designated as “low” (**Fig. 5B, Fig. S6D**). In this repeat RNA-ISH dataset, the HERVK and HERVH RNA species quantities were well correlated with LINE1 (**Fig. 5C**), however, the HSATII repeat was not linearly correlated as we have previously described across other epithelial cancers (9,44). The relationship between expression of LINE1 and overall survival (OS) was assessed using Kaplan Meier analysis and log-rank test to determine significance (**Fig. 5D**). High LINE1 expression was associated with worsened overall survival (median OS 2.04 vs. undefined years, p = 0.01).

**Figure 5.**
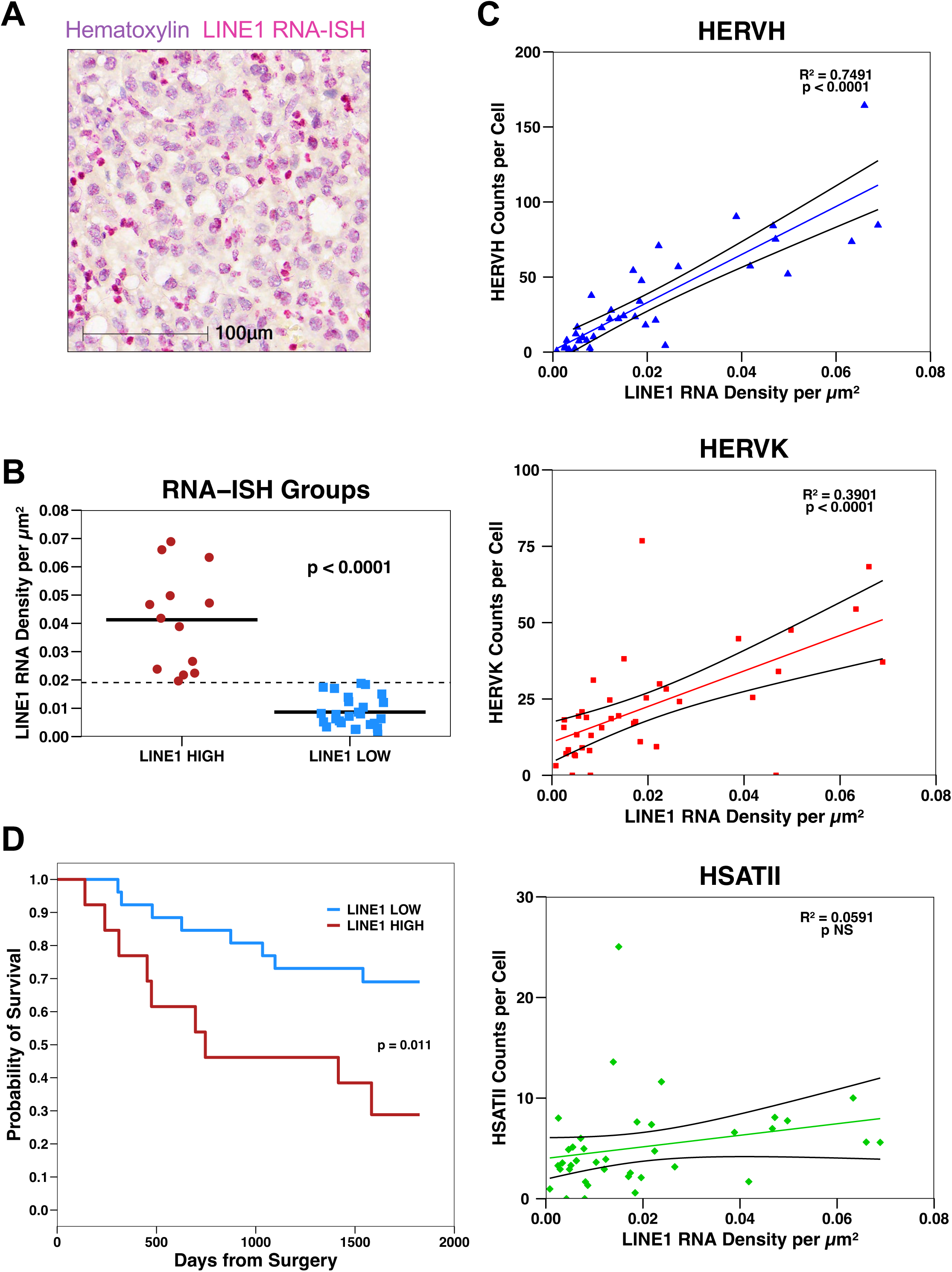
Validation of LINE1 expression by RNA-ISH. (**A**) Representative RNA-ISH image showing LINE1 expression. (**B**) Jitter plot depicting the definition of LINE1 high and low patient groupings in the RNA-ISH dataset. The dashed line represents the upper tercile for LINE1 counts per µm^2^ across all patients. The crossbars represent means. P-value computed by Wilcoxon Rank Sum Test. (**C**) Association of LINE1 RNA-ISH quantification with HERVH, HERVK, and HSATII RNA-ISH quantification. P-values computed by Wald test. (**D**) Kaplan-Meier overall survival analysis for LINE1 high patients versus LINE1 low patients (p = 0.01 by log-rank test).

## DISCUSSION

The development, progression, and response to systemic therapy of hepatocellular carcinoma is influenced by complex interactions between cancer cells and the stromal/immune tumor microenvironment. In this study, we utilized a single cell resolution spatial transcriptomic profiling technology to characterize how the HCC tumor microenvironment (TME) varies in a manner dependent on cancer cell expression of the LINE1 repeat RNA. The spatial molecular imaging platform and downstream computational analytic pipeline enabled robust gene expression profiling of thousands of single cells per sample with retained spatial context. Comprehensive single cell spatial analysis including stromal cells and lymphoid cells provide a high resolution map of the HCC TME. These data will unlock new therapeutic opportunities T cells and endothelial cells, which have shown to be the most pertinent druggable targets in the HCC TME (45).

Given the prominent role of the LINE1 retrotransposon in cancer (46), we chose to focus first on LINE1 expression as a marker of aggressive cancer phenotype. We found that LINE1 expression had elevated expression of repeat RNAs HSATII and HERVK, consistent with coordinate de-repression of these critical repeat elements observed other cancer types (47). This was validated by RNA-ISH, but we found that HSATII was not linearly correlated with LINE1 expression consistent with distinct regulation of diverse repeat sequences we have observed in ovarian, pancreatic, and colorectal cancers (9,44). LINE1-high tumors and single cells exhibited an aggressive stem-like (POU5F1, TWIST2 (48)) and immune suppressive phenotype (PDCD1); LINE1-low tumor and single cell expression profiles showed less aggressive, more well-differentiated hepatocyte features (SAA1/2, APOA1). Leveraging the retained spatial data in combination with single cell expression profiling allowed detailed analysis of the microscopic cellular neighborhoods associated with LINE1 expression. We noted that LINE1 cancer cell expression was correlated with significant differences in the TME. Whereas LINE1 high cells tended to surround themselves with additional LINE1 high cells, LINE1 low cells associated with other LINE1 low cells and immune cell aggregates. Other groups have shown that “tertiary lymphoid structures” (TLSs) in HCC (49) and “immunity hubs” in non-small cell lung cancer (41) correlate with effective anti-tumor immune responses. Our findings suggest that LINE1 overexpression may serve as a biomarker for disorganized immune responses to cancer and imply that disrupting LINE1 effects may augment anti-tumor immunity.

We validated our findings with an extended cohort of RNA-ISH specimens, confirming coordinate expression of LINE1 with other repeat RNA elements. We also noted that high expression of LINE1 in HCC tissue specimens was associated with poorer survival, consistent with prior work in other tumor types (50,51) and in hepatitis B driven HCC (33). Other analyses have identified hypomethylation of LINE1 as adverse prognostic factors in various HCC subsets (27–30). Our study is the first to identify LINE1 transcript expression as a potential biomarker of poor prognosis in HCC. Given the observation that robust pre-resection immune infiltrates correlate with good surgical outcomes in HCC (49) and other (52) cancers, LINE1 associated immune disarray may be driver of this effect.

Our study implies that LINE1 – and perhaps additional co-regulated repeat RNAs such as HSATII and HERVK – may represent potential novel therapeutic targets in HCC. HERVK is considered the most translationally active of the endogenous retroviruses, encoding for proviral Gag, Pro, Pol, and Env proteins (53). Specific targeting of HERVK proteins with monoclonal antibodies (54), HERVK reactive cytotoxic T-cells derived from patient serum (55), and CAR T-cells (56) have demonstrated anti-tumor effects in preclinical models. LINE1 encodes for ORF1, an RNA-binding protein, and ORF2, which has endonuclease and reverse transcriptase activity. Circulating LINE1-ORF1 has been proposed as a pan-cancer (including HCC) diagnostic marker (57), and LINE1-ORF1p has been explicitly identified as a potential therapeutic target in pancreatic cancer. These protein products of LINE1 and other repeat RNA elements may similarly be targetable in HCC.

The limitations of this study include the small sample size and the retrospective nature of the analyses. The tissue samples were operative samples from systemic therapy-naïve patients who underwent surgical resection or transplantation for management of their HCC, and therefore patients with unresectable or metastatic disease who may have different tumor biology at baseline or in response to systemic therapy regimens were not represented in the cohort. Additionally, while our spatial transcriptomic platform leveraged expression of thousands of genes, the majority of the (coding and non-coding) transcriptome was not assayed. Finally, while distinct tumor immune profiles were found correlate with LINE1 RNA expression, additional mechanistic experiments are needed to further assess potential causal relationships.

In summary, our study is the first apply single cell spatial transcriptomic technologies simultaneously to coding and non-coding RNA elements in HCC. We identified expression of the LINE1 retrotransposon as a potential adverse RNA prognostic biomarker and marker of tumor de-differentiation/stemness and disarrayed immune infiltrates. This work expands upon the existing literature describing the interplay between the coding and non-coding transcriptome in cancer and lays the groundwork for the development of novel biomarkers and mechanistic hypotheses to test in subsequent investigations. Ultimately, our data may motivate the development of additional therapeutic strategies to augment anti-tumor immunity and improve clinical outcomes for patients with HCC.

### Materials Availability

All RNA-ISH probes and sequences are available at ACD-Biotechne and custom CosMx probes from Bruker.

### Data and Code Availability

All code and statistical packages are detailed in the methods and supplemental methods and will be made available upon request. All images and primary data will be made available upon request. Spatial transcriptomic expression matrices will be available upon request.

## AUTHORS’ DISCLOSURES

JWF has received consulting fees from Eisai, Foundation Medicine, Guardant Health, Genentech, and Servier. JWF has received personal research funding from Genentech and institutional research funding from Abbvie, Alnylam, Genentech, Iterion Therapeutics, and Omega Therapeutics. None of these competing interests are related to this work. DTT has received consulting fees from ROME Therapeutics, Sonata Therapeutics, Leica Biosystems Imaging, PanTher Therapeutics, 65 Therapeutics, and abrdn. DTT is a founder and has equity in ROME Therapeutics, PanTher Therapeutics and TellBio, Inc., which is not related to this work. DTT is on the advisory board with equity for ImproveBio, Inc. and 65 Therapeutics. DTT has received honoraria from Astellas, AstraZeneca, Moderna, and Ikena Oncology that are not related to this work. DTT receives research support from ACD-Biotechne, AVA LifeScience GmbH, Incyte Pharmaceuticals, Sanofi, and Astellas which was not used in this work. DTT’s interests were reviewed and are managed by Mass General Brigham in accordance with their conflict of interest policies.

## AUTHOR CONTRIBUTIONS

Conceptualization AKC, CN, BKP, JWF, DTT

Formal Analysis CN, AKC, ZG, YS, KHX, AC, LTN, MJA, JWF, DTT

Investigation by AKC, CN, BKP, AP, ZG, MJE, ERL, YS, KHX, LTN, MJA, JWF, DTT

Methodology by AKC, CN, ZG, YS, KHX, LTN, MJA, JWF, DTT

Resources by MJE, CRF, VD, LTN, JWF, DTT

Writing AKC, CN, BKP, MJA, JWF, DTT

Visualization by AKC, CN, KHX, LTN, DTT

Supervision by LTN, MJA, JWF, DTT

Project Administration MJA, JWF, DTT

Funding Acquisition AKC, JWF, DTT

## ACKNOWLEDGMENTS

We thank Angelique Gilbert and Danielle Bestoso for laboratory and administrative support. We are thankful for the patients that provided tumor material for this project. This work was supported by research funding from the National Institutes of Health R01CA240924 (DTT), U01CA228963 (DTT), 3R01CA240924-04S1 (AC), K08CA263551 (JWF), ACD-Biotechne (DTT), and the Krantz Family Center for Cancer Research Quantum Grant (DTT).

## Notes

### Summary of Updates

This revised manuscript has changed significantly with addition of Single Molecule Imaging data (CosMx) in HCC with addition of authors (Cole Nawrocki, Zhoubou Guo) to reflect changes of content. The figures are significantly different and some data was removed as it is no longer relevant for this particular manuscript.

