## Supplementary Text for "Single Cell Spatial Transcriptomic Profiling Identifies a LINE1 Associated Disarrayed Immune Microenvironment in Hepatocellular Carcinoma"

**SUPPLEMENTAL METHODS**

**CosMx Data QC and Processing**

Each field of view (FOV) was reviewed, and *Cellpose* (1) cell segmentation was performed, using NanoString’s preset configurations A or E for the “cyto2” model, based on which appeared to perform best. Next, cells that did not exhibit the following characteristics were discarded: total RNA counts greater than 20 and less than 2000, cell area greater than 25 µm^2^ and less than 1000 µm^2^, proportion of total counts from negative control probes less than 0.1, and complexity score less than 2. Complexity score was defined as log10[(total RNA counts) / (positive RNA genes)]. Thus, cells with 100 times as many counts as positive genes were considered erroneous and were discarded. After filtering poor quality cells, FOVs with more than 25% of their cells deemed poor quality were themselves entirely discarded to preserve general contiguity of the tissue structure for downstream spatial analysis. Core 22 was removed from TMA2 due to high levels of necrosis and outlier-level expression of spike-in probes. 91.6% and 83.6% of cells from TMA1 and TMA2 were retained, respectively.

Cells from each TMA were then processed separately, using *Seurat* (2). First, expression was normalized to log_2_(counts per thousand + 1) and z-score scaled for all genes. This scaled expression matrix was used to generate 50 principal components, 25 of which were used to generate a UMAP embedding for each TMA. After cell-typing was performed for each TMA, all data were merged. This merged data set was then subject to the same processing scheme. No batch effect was observed between the two TMAs.

**CosMx Data Cell-typing**

First, tumor cells and erythrocytes were identified manually for each TMA. To identify tumor cells, a tumor meta score was generated for each cell, by inputting the following genes into *Seurat*’s *AddModuleScore* function: APOA1, GC, VTN, ARG1, FGG, and GPC3. Louvain clustering was performed with resolution parameters of 0.5 and 0.7 for TMA1 and TMA2 respectively. By visualizing the resulting clusters and the tumor meta score on the UMAP embedding, tumor clusters were found. To identify erythrocytes, all genes with 3 blue bits present in their reporter probe barcode were passed to *AddModuleScore* to produce an autofluorescence meta score. As with tumor cells, using Louvain clustering and visualization, erythrocyte clusters were found.

Next, using liver (3) and immune-oncology reference profiles, supervised clustering was performed using *InSituType* (Danaher et al, BioRvix 2022)*.* on the remaining cells to identify final cell types. Spatial nearest neighbor expression produced from each cell’s 50 nearest spatial neighbors was converted to 10 principal components. These components, along with each cell’s immunofluorescent staining data and cell area was passed to *InSituType* as cohort data to improve clustering. When clustering with the immune-oncology profile, genes that were likely sources of spatial contamination were excluded.

**CosMx Data Cell-typing Validation**

After cell-typing, the data from both TMAs were merged, and marker genes were found for each cell type cluster, using two methods. First, an algorithmic approach (4) identified genes that are uniquely highly expressed in each cluster. First, the mean counts per cell was found for each gene in each cluster. For each cluster, all genes with a counts per cell value greater than both 0.1 and the second-largest counts per cell value in any other cluster were identified. These genes were then ranked according to how much larger their counts per cell value was in each cluster, compared to their next largest counts per cell value in any other cluster. Finally, for each cluster, the top 1-5 ranked genes were selected as marker genes for visualization.

Second, a modeling approach identified significant differentially expressed genes that characterize each cluster. Specifically, after pseudo-bulking, the *presto-GLMM* R package was used to fit a Poisson GLMM for each gene’s counts with the following formula: y_g ~ 1 + (1|celltype) + (1|celltype:patient) + (1|patient) + offset(logUMI). Linear contrasts representing each cluster versus the rest were tested for each gene’s model to find logFC values and corresponding p-values via a Wald test. These p-values were FDR-corrected. For each cluster, genes with logFC greater than 1.5 and adjusted p-value less than 0.05 were considered marker genes. Review of each cluster’s marker genes demonstrated that high fidelity annotations had been made.

**CosMx Data Differential Expression Analysis**

To identify LINE1-ORF1 high patients, the total LINE1-ORF1 counts in all tumor cells was quantified for each patient. This measure was converted to counts per million (CPM), and the patients in the upper tercile for LINE1-ORF1 CPM were labeled “high,” while the remaining patients were labeled “low.” Next, each gene’s counts in tumor, endothelial, and macrophage cells were modeled with a Negative Binomial GLMM using the *NEBULA* (5) package and the following formula: y_g ~ line1orf1_group + celltype + line1orf1_group:celltype + (1|patient) + offset(logUMI). A linear contrast representing “high” tumor cells versus “low” tumor cells was tested for the model with an asymptotic Wald test for each gene, and the resulting p-values were FDR-adjusted. Genes that were likely sources of spatial contamination were not modeled or tested. This same scheme was applied to endothelial cells as well.

To identify LINE1-ORF1 high tumor cells, within each patient, cells in the upper tercile for LINE1-ORF1 normalized expression were labeled “high” and the remaining cells were labeled “low.” Tumor cells in each group were pseudo-bulked to form “high” and “low” paired tumor samples for each patient. Next, each gene’s counts in tumor samples were modeled with the *DESeq2* (6) package using the following formula: y_g ~ line1orf1_group + patient. During this process, size factors were calculated for each sample, using the upper quartile method from *edgeR* (7). Finally, a linear contrast representing “high” tumor cells versus “low” tumor cells was tested with a likelihood ratio test. For this, we set parameters fitType = “glmGamPoi” and reduced = ~patient.

**CosMx Data Gene Set Enrichment Analysis**

Using our pseudo-bulk based differential expression analysis results, we omitted spike-ins and likely contaminated genes, then ranked all remaining genes in descending order of their likelihood ratio test statistic multiplied by their logFC sign. This pre-ranked gene list was passed to the GSEA function from the *clusterProfiler* (8) package. We tested all gene sets in the C8 and GO:MF categories from MSigDB (9).

**CosMx Data Neighborhood Enrichment Analysis**

We conducted neighborhood enrichment analysis using methods inspired by Varrone et al (10). Within each FOV, we computed a Delaunay tessellation. For each cell, which we refer to as the index cell, we defined the neighbor cells to be the set of directly adjacent cells on this tessellation. We combined FOV-wise neighborhood definitions to form a global neighborhood graph, trimming all graph edges with an associated distance value greater than 100 µm. Thus, by this scheme, for a cell to be a neighbor of a given index cell, the cell must exist in the same FOV as the index; the cell must be directly adjacent to index, with no cells between them; and the cell must be within 100 µm of the index.

After partitioning tumor cells into LINE1-ORF1 high and low subsets in a patient-wise fashion, as previously described, we aimed to assess high and low tumor cell neighborhoods for comparison. To accomplish this, we did the following for each cell-type, including LINE1 high and low tumor cells: 1) In a patient-wise fashion, for a given cell type, we treated all cells of that type as index cells. 2) We found the proportions of neighbors to all index cells that belonged to each cell type identity. 3) We defined these proportions as the observed proportions of neighbors to the index cell type. 4) We defined the expected proportions of neighbors to the index cell type analytically. The expected proportion of neighbors of a specific cell type identity, which we will denote as cell type X, was defined as the total number of neighborhood graph edges connecting to cells of type X, divided by the total number of edges in the graph. 5) Adding a small pseudo-count of 0.0001 to the observed and expected proportion values, then taking log2 of the (observed/expected) ratio defined the log2 enrichment ratio (log2ER), a measure of colocalization. 6) We compared log2ER profiles between “high” and “low” tumor index cells, using a patient-paired Wilcoxon Rank Sum Test.

Second, we aimed to compare the cell co-localization patterns in LINE1-ORF1 high and low patients. To accomplish this, we first split the data set by patient-level LINE1-ORF1 group. Just as previously described, we computed neighborhood graphs for each partition of the data set. Without splitting by individual patient, we computed log2ER values for each. To assess significance for differences in log2ER values across the two partitions, we used simulation. We randomly permuted the patient grouping labels 1000 times. During each permutation, we computed the difference between the log2ER values in the “high” group and the log2ER values in the “low” group. Comparing a given observed difference to the simulated differences allowed us to calculate empirical p-values. To test for a difference in the overall immune cell colocalization pattern across the “high” and “low” patient groups, we defined the sum of all pairwise log2ER values for all immune cell types as the immune colocalization score. Next, we found the difference in this score across the “high” and “low” patient groups, which we defined as the observed value. Recording this same metric during each of the 1000 simulations detailed above allowed us to define an empirical null distribution, which we used to calculate an empirical p-value.

**Statistical survival analysis**

Kaplan Meier analysis with log-rank test for significance was performed using Graphpad Prism (v10).

**Data and Code Availability**

All data will be provided upon request. Analysis code is available at <https://github.com/ccnawrocki/hcc-tma-cosmx-project>.

**SUPPLEMENTAL FIGURES**

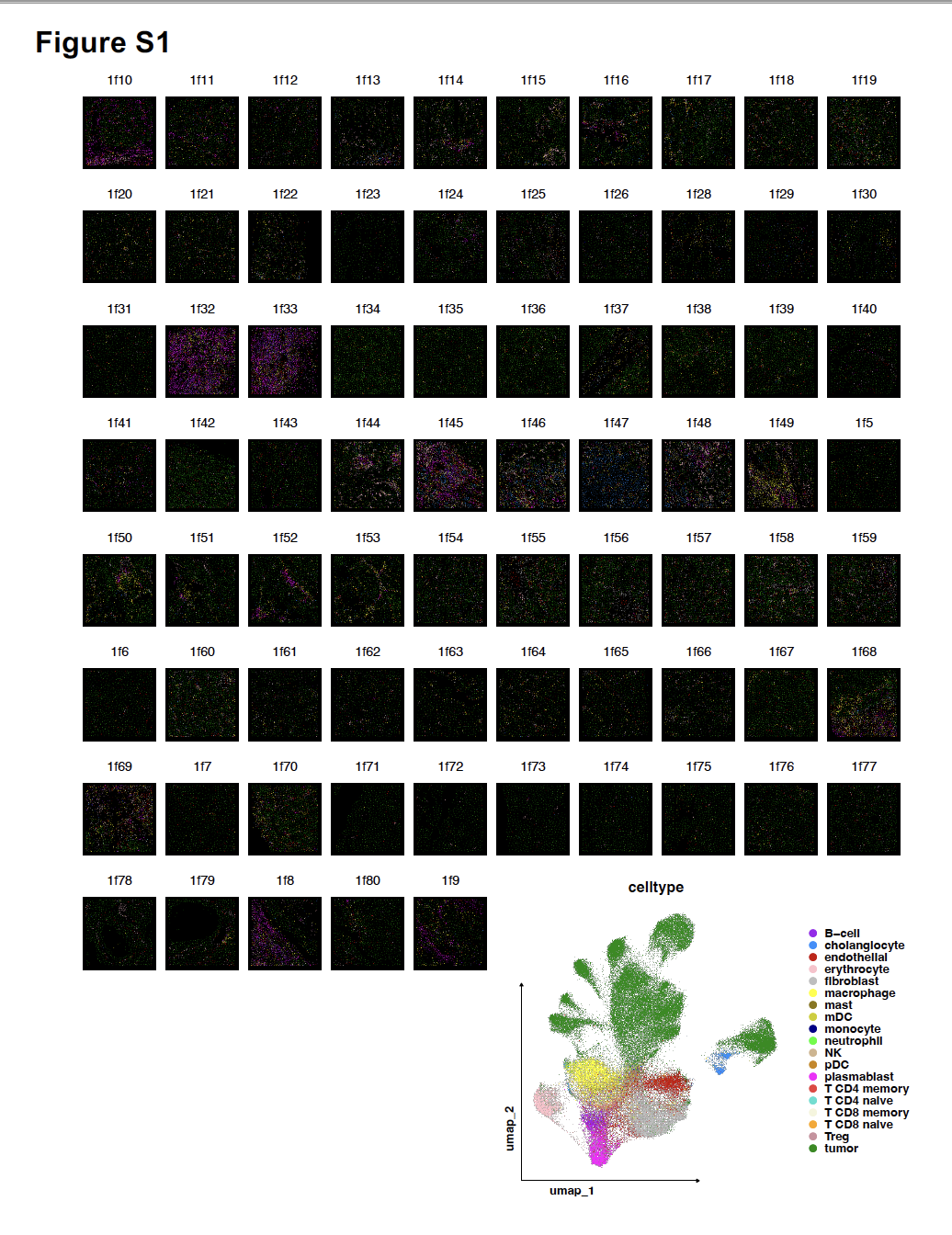

**Figure S1. Spatial mapping of cell type annotations for all fields of view (FOVs) on TMA1.** Main panel depicts FOV-wise spatial maps (see Figure S3A for context). Bottom panel shows a UMAP embedding with cells pseudo-colored by cell type annotation for all cells on TMA1.

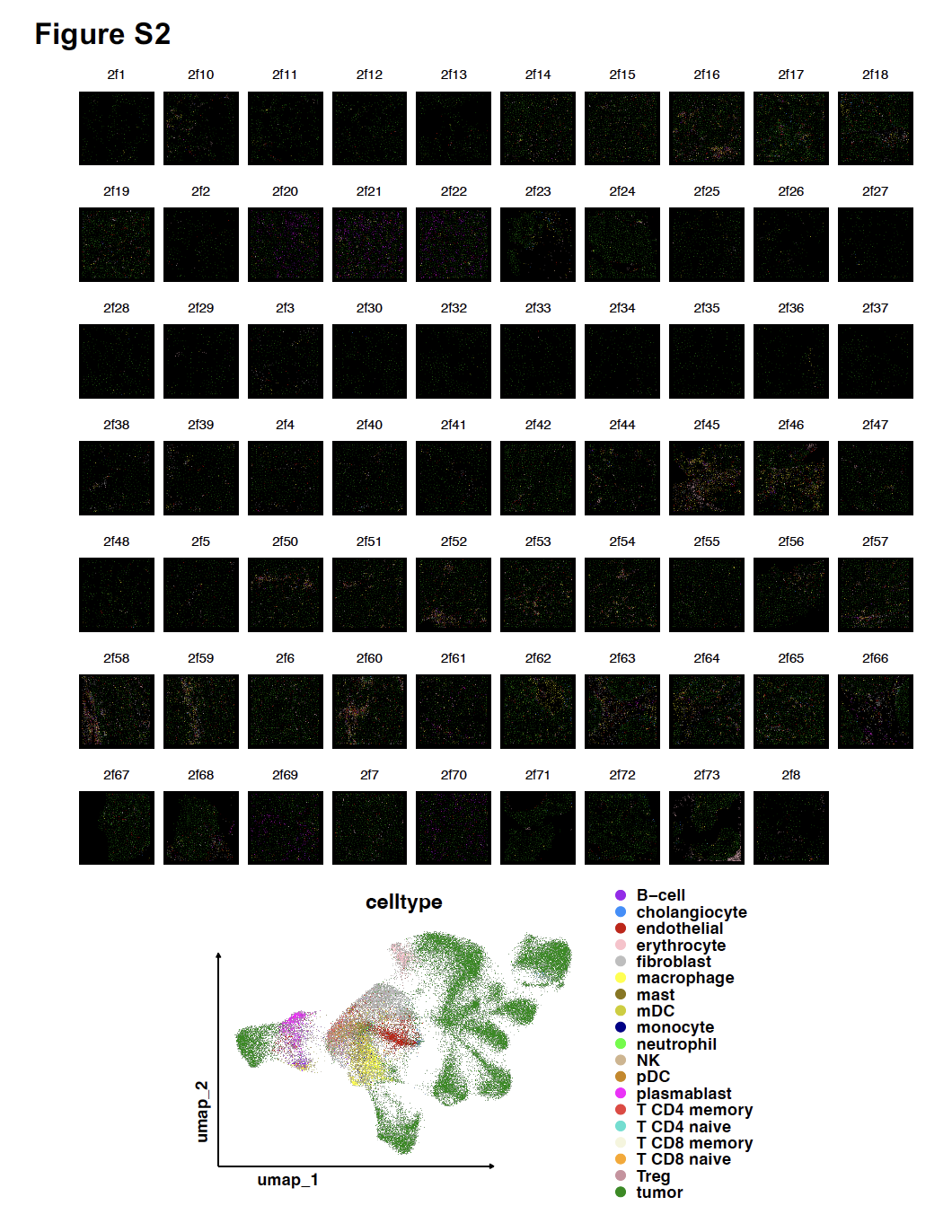

**Figure S2. Spatial mapping of cell type annotations for all fields of view (FOVs) on TMA2.** Main panel depicts FOV-wise spatial maps (see Figure S3B for context). Bottom panel shows a UMAP embedding with cells pseudo-colored by cell type annotation for all cells on TMA2.

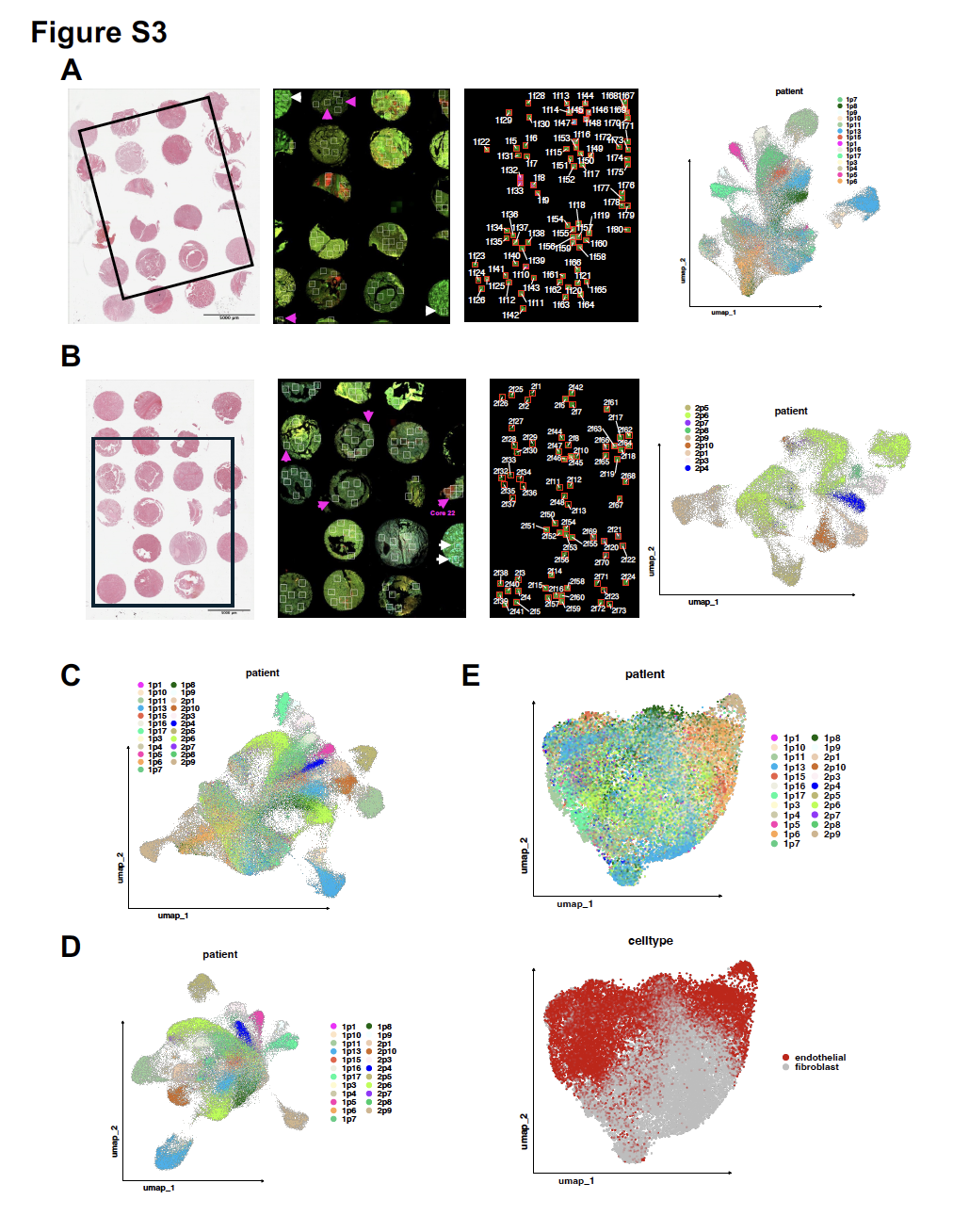

**Figure S3. Acquisition of spatial molecular imaging (SMI) data for two HCC FFPE TMAs allows for quantitative isolation of specific cell type subsets.** (**A**) Left: H&E staining for TMA1 with the region selected for SMI enclosed in black. Second from left: immunofluorescent staining for the selected region in S3A. Each white rectangle depicts a field of view (FOV), within which SMI was conducted. Pink arrows indicate FOVs filtered due to QC. White arrows indicate kidney control FOVs. Staining matches that in Figure 1B. Second from right: spatial map of relative TMA1 FOV locations with cells pseudo-colored by cell type annotation inside each FOV. Right: UMAP embedding for TMA1 with cells pseudo-colored by patient identity. (**B**) Replication of S3A, for TMA2. Note that core 22 was filtered entirely due to QC. (**C**) UMAP embedding for the complete SMI dataset with cells pseudo-colored by patient identity. (**D**) UMAP embedding for all tumor cells identified in the SMI dataset with cells pseudo-colored by patient identity. (**E**) UMAP embedding for all stromal cells identified in the SMI dataset with cells pseudo-colored by patient identity (top) and by cell type annotation (bottom).

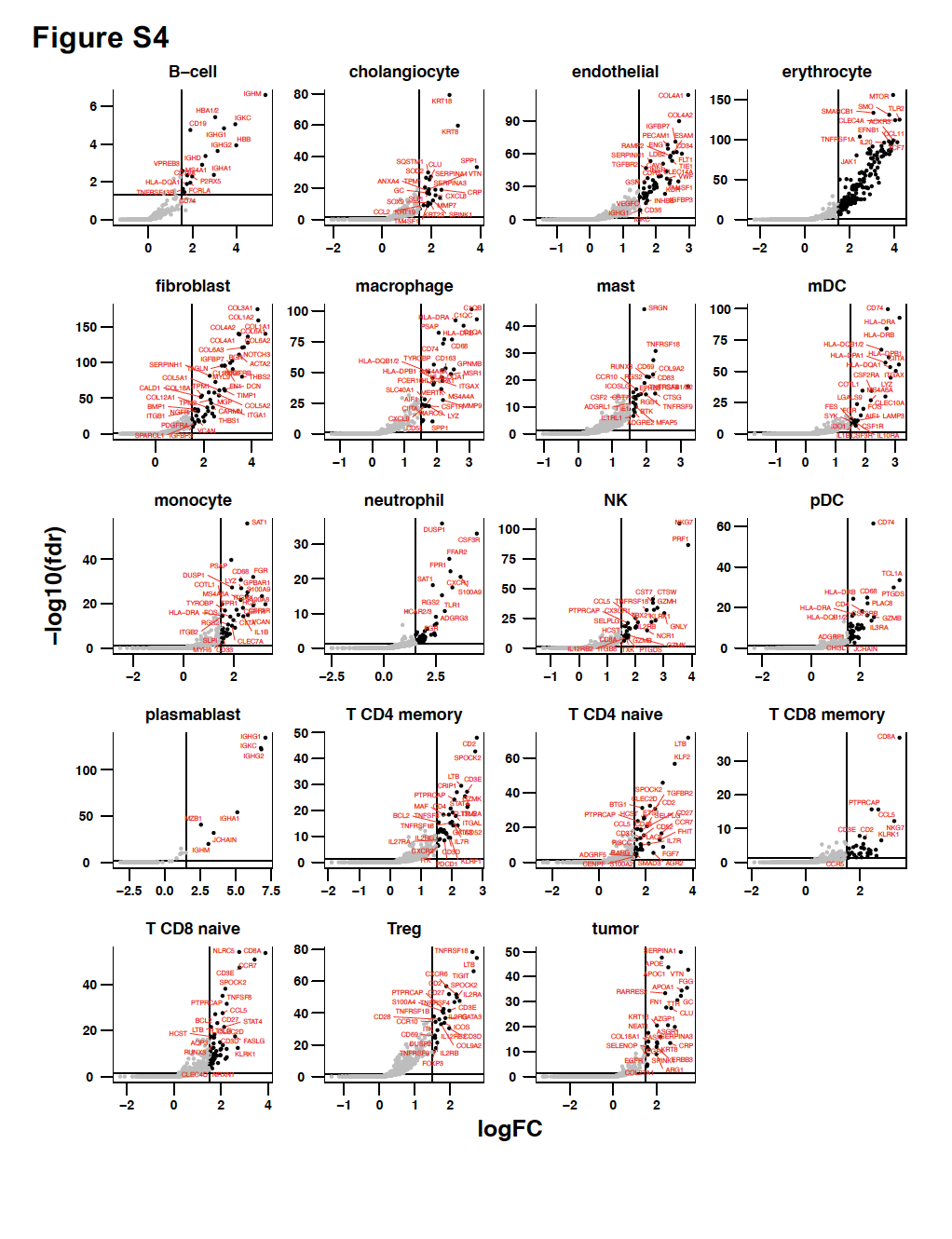

**Figure S4. Validation of cell-typing with differentially expressed marker genes for cell type clusters.** One-sided volcano plots depicting significant differentially expressed positive marker genes for each cell type cluster (see methods). Each cluster was compared to all other clusters. P-values computed by Wald test and FDR-adjusted. Significance threshold = 0.05. Effect size threshold = logFC value of 1.5.

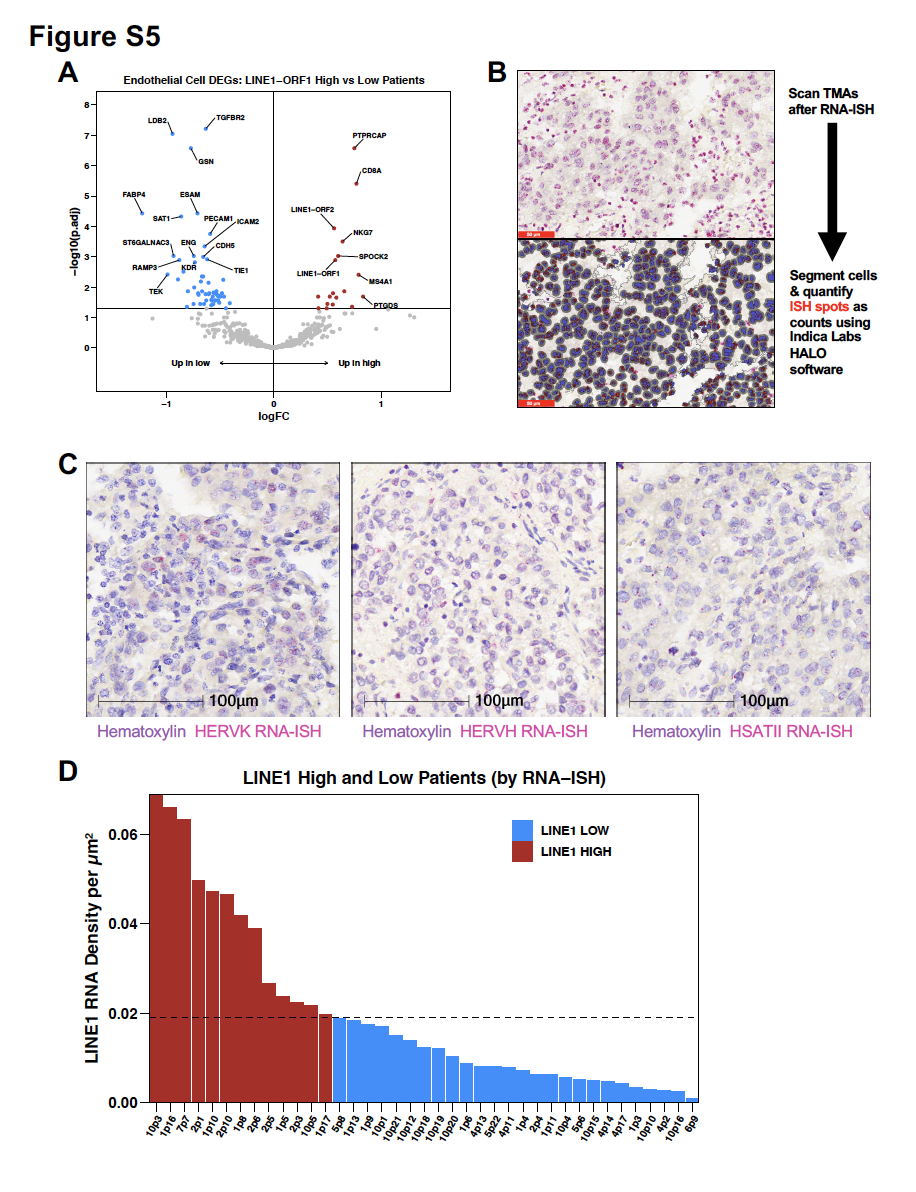

**Figure S5. Continuation of differential expression analysis in endothelial cells and extended information for RNA-ISH validation.** (**A**) Volcano plot summarizing differential expression analysis comparing endothelial cells in LINE1-ORF1 high patients and LINE1-ORF1 low patients, as defined in Figure 3A. P-values are FDR-adjusted. Significance threshold = 0.05. (**B**) Schema of RNA-ISH data collection and quantification workflow. For each cell, the nucleus is pseudo-colored purple, the cytoplasm is pseudo-colored grey, and confidently identified RNA-ISH spots are pseudo-colored red (bottom). (**C**) Representative images of HERVK, HERVH, and HSATII RNA-ISH staining (left to right). (**D**) Bar plot depicting the definition of LINE1 HIGH and LOW patient groupings in the RNA-ISH dataset. The dashed line represents the upper tercile for LINE1 RNA count density (per µm^2^) across all patients.

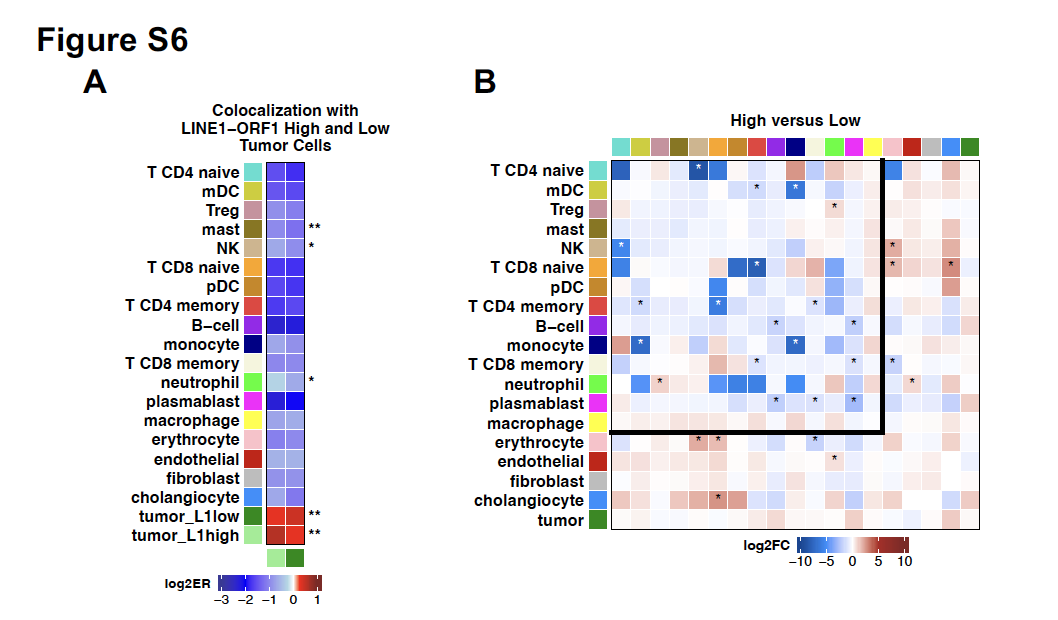

**Figure S6. Extended information for neighborhood enrichment analysis.** (**A**) Differences in LINE1-ORF1 high and low tumor cells’ colocalization patterns with themselves, one another, and other cell types across the complete SMI dataset. ER = enrichment ratio (see methods). P-values computed by paired Wilcoxon Rank Sum Test and FDR-adjusted. * = nominal p-value < 0.05. ** = adjusted p-value < 0.05. (**B**) Observed differences in all pairwise cell type colocalizations between LINE1-ORF1 high patients and LINE1-ORF1 low patients in the SMI dataset (See Figure 4C). Differences were computed as high minus low. P-values computed empirically. * = nominal p-value < 0.05.

**SUPPLEMENTAL TABLE TITLES**

**Table S1. List of genes targeted by spatial transcriptomic oligonucleotide probe set**.

**Table S2. Differential expression results for LINE1-high versus LINE1-low tumor cores.**

**Table S3. Differential expression results for LINE1-high versus LINE1-low single cells**.
